# Biochemical and functional characterization of the p.A165T missense variant of mitochondrial amidoxime-reducing component 1 in HepG2 cells

**DOI:** 10.1101/2023.10.18.562989

**Authors:** Wangfang Hou, Christian Watson, Ted Cecconie, Menaka N. Bolaki, Jennifer J. Brady, Quinn Lu, Gregory J. Gatto, Tovah A. Day

## Abstract

Recent genome-wide association studies have identified a missense variant p.A165T in mitochondrial amidoxime-reducing component 1 (mARC1) that is strongly associated with protection from all-cause cirrhosis and improved prognosis in nonalcoholic steatohepatitis (NASH). The precise mechanism of this protective effect is unknown. Substitution of alanine 165 with threonine is predicted to affect mARC1 protein stability and to have deleterious effects on its function. To investigate the mechanism, we have generated a knock-in mutant mARC1 A165T in human hepatoma HepG2 cells, enabling characterization of protein subcellular distribution, stability, and biochemical functions of the mARC1 mutant protein expressed from its endogenous locus. Compared to wild-type (WT) mARC1, we found that the A165T mutant exhibits significant mislocalization outside of its traditional location anchored in the mitochondrial outer membrane and reduces protein stability, resulting in lower basal levels. We evaluated the involvement of the ubiquitin proteasome system in mARC1 A165T degradation and observed increased ubiquitination and faster degradation of the A165T variant. In addition, we have shown that HepG2 cells carrying the *MTARC1* p.A165T variant exhibit lower N-reductive activity on exogenously-added amidoxime substrates *in vitro*. The data from these biochemical and functional assays suggest a mechanism by which the *MTARC1* p.A165T variant abrogates enzyme function which may contribute to its protective effect in liver disease.

## Introduction

Nonalcoholic fatty liver disease (NAFLD) is a major cause of chronic liver disease. NAFLD affects over 64 million people in the US and approximately 2 billion people worldwide (1,2). About 10% of NAFLD patients progress to the most serious form, nonalcoholic steatohepatitis (NASH). Cirrhosis, a severe complication of NASH and other chronic liver diseases, is the 11^th^ leading cause of death globally. It accounts for ∼35,000 deaths annually in the US and represents a major public health burden (3-7). Despite this, there are currently no FDA-approved therapies for NASH (8).

Genome-wide association studies (GWAS) are a powerful tool to uncover associations between genetic variants and risk of disease; recently, they have also become an important strategy to identify new therapeutic targets (9,10). Several recent studies applied GWAS to NASH and identified the missense mitochondrial amidoxime-reducing component 1 (mARC1) variant p.A165T as protective from both alcoholic and nonalcoholic cirrhosis (1,10,11). *MTARC1* p.A165T interfered with lipid metabolism (12,13). It has also been associated with lower liver enzymes, lower total cholesterol level (1,10), reduced severity of NAFLD and reduced liver-related mortality (1,12). This protective effect of p.A165T is very similar to a rare nonsense mutation p.R200Ter, which completely lacks the catalytic domain at the C-terminus of the protein (10). A subsequent study showed that *MTARC1* p.A165T variant was implicated in downregulation of the hepatic fibrotic pathway and connected with lower grade of hepatic steatosis in children with NAFLD (13).

mARC1 protein (encoded by the *MTARC1* gene) is one of four molybdenum (Mo)-containing enzymes in the human genome (14). It localizes to the outer mitochondrial membrane and is predominantly expressed in liver and adipose tissue (15,16). The C-terminal domain is exposed to the cytosol where it binds to a molybdenum cofactor (Moco) and functions as the catalytic core. The N-terminal domain contains a mitochondrial targeting sequence and a hydrophobic domain which anchors the protein on the outer mitochondrial membrane (15). mARC1 forms a complex with cytochrome *b*5 reductase 3 (CYBR3) and cytochrome *b*5 type B (CYB5B), from which Moco receives electrons from NADH to enable mARC1 to reduce a large variety of N-oxygenated substrates including nitrite, N-hydroxylated nucleobases, amidoxime prodrugs and physiological substrate *N*^ω^-hydroxy-L-arginine (NOHA) (17-22). Based on its enzymatic activity, mARC1 is hypothesized to play roles in cellular detoxification, L-arginine metabolism, nitric oxide (NO) biosynthesis and drug metabolism (20,23,24). NO affects mitochondrial biogenesis and inhibits mitochondrial respiration, thereby influencing mitochondrial functions both physiologically and pathologically (25-28). Mitochondrial dysfunction leads to increased formation of reactive oxygen species (ROS) and lipid peroxidation, which both contribute to NASH (29). These clinical observations, coupled with the GWAS finding (1,10,11) strongly suggest a role for mARC1 in NASH, however, the mechanistic details remain poorly understood.

A total of 27 nonsynonymous single nucleotide polymorphisms (SNPs) have been described for *MTARC1* (30). Among these, c.493A>G (rs2642438, resulting in p.A165T) has a minor allele frequency of 0.25 in the human population (31). *In silico* modeling predicted that *MTARC1* p.A165T variant would have lower stability and altered molybdenum binding which would abrogate mARC1 enzymatic function (13). However, purified, recombinant mARC1 A165T exhibited similar kinetic parameters to WT when reducing benzamidoxime (30) and sulfamethoxazole hydroxylamine (17), two N-hydroxylated substrates. These observations suggest that A165T does not abrogate *in vitro* binding of molybdenum since Moco binding is essential for N-reductive function of mARC1 (30). Despite the lack of difference in recombinant enzyme turnover, GWAS of *MTARC1* p.A165T shows remarkably broad protection from liver fibrosis, steatosis and cirrhosis and a benefit in NASH, indicating that elucidation of a protective mechanism in a cellular context will yield important insights into this modulator of liver diseases. Our exploration in this study provides biochemical and functional explanations for the protection effect of this variant using a cellular model.

## Results

### mARC1 A165T protein is mislocalized in HepG2 cells

mARC1 is a mitochondrial protein localized to the outer mitochondrial membrane. To investigate whether single amino acid changes in mARC1 variants affected protein subcellular localization, we imaged mARC1 protein and mitochondria in HepG2 cells. In cells expressing the A165T variant, more diffuse staining of mARC1 protein was observed when compared to WT and C273A, a catalytically dead mutant (Fig. 1A). Colocalization analysis of mARC1 staining and mitochondrial staining revealed that the A165T variant co-localize with mitochondria significantly less than WT (Fig. 1A). This was quantitated as FITC intensity of mARC1 protein for mitochondrial puncta and background for an average from over 6,000 cells. The average corrected FITC spot intensity of mitochondrial puncta of A165T was significantly lower than that of WT or C273A but indistinguishable from the background (Fig. 1B).

**Figure 1.**
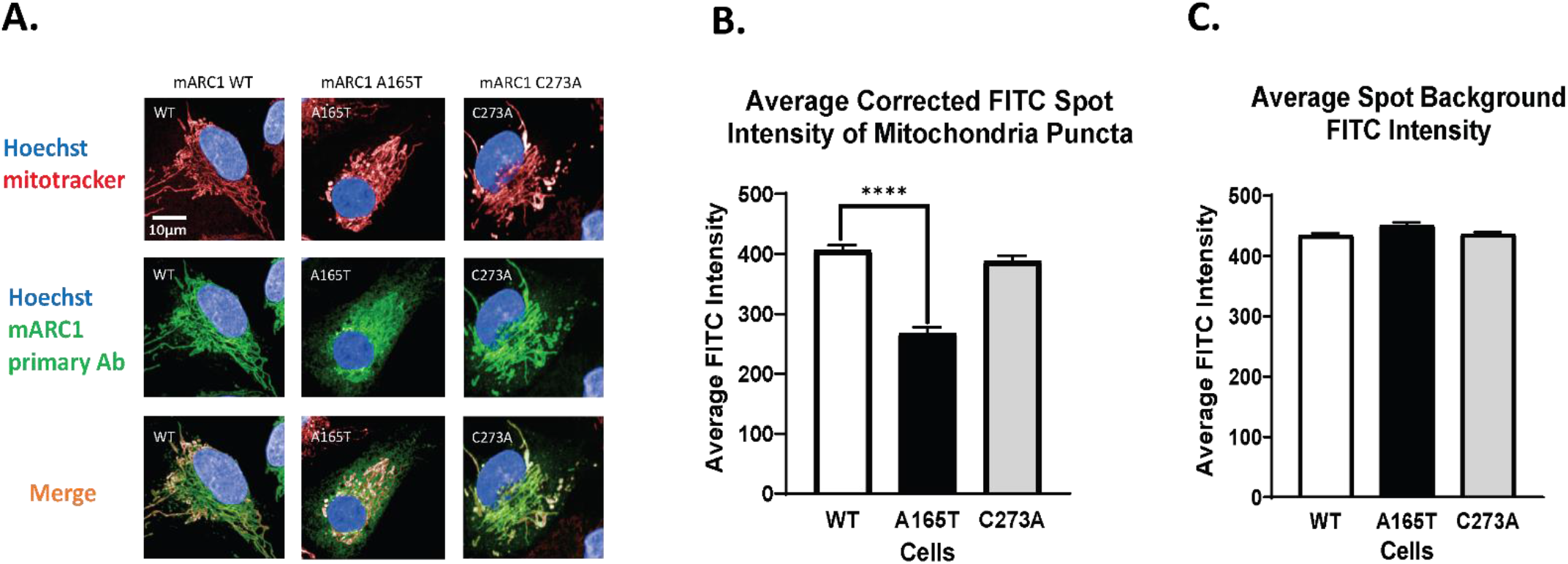
Subcellular localization of mARC1 variants and comparison of their level on mitochondria. A. Imaging data for colocalization of mARC1 and mitochondria; B. Average corrected FITC (mARC1) spot intensity of mitochondria puncta and average spot background FITC intensity. Data represents an average from 6 wells and each well has >1000 cells well analyzed (**** P<0.0001).

### A165T destabilizes mARC1

To further characterize the effect of A165T on mARC1, we examined the endogenous level of mARC1 expression in CRISPR-edited HepG2 cells expressing different variants. mRNA and protein level of mARC1 were determined for the three HepG2 cell lines carrying different *MTARC1* genotypes including WT, A165T and C273A. There was no significant difference in the levels of mRNA between A165T and WT (Fig. 2A). The negative control, HepG2 mARC1 KO cells deficient in mARC1, exhibited dramatic (>90%) loss of mARC1 mRNA relative to WT (Fig. 2A).

**Figure 2.**
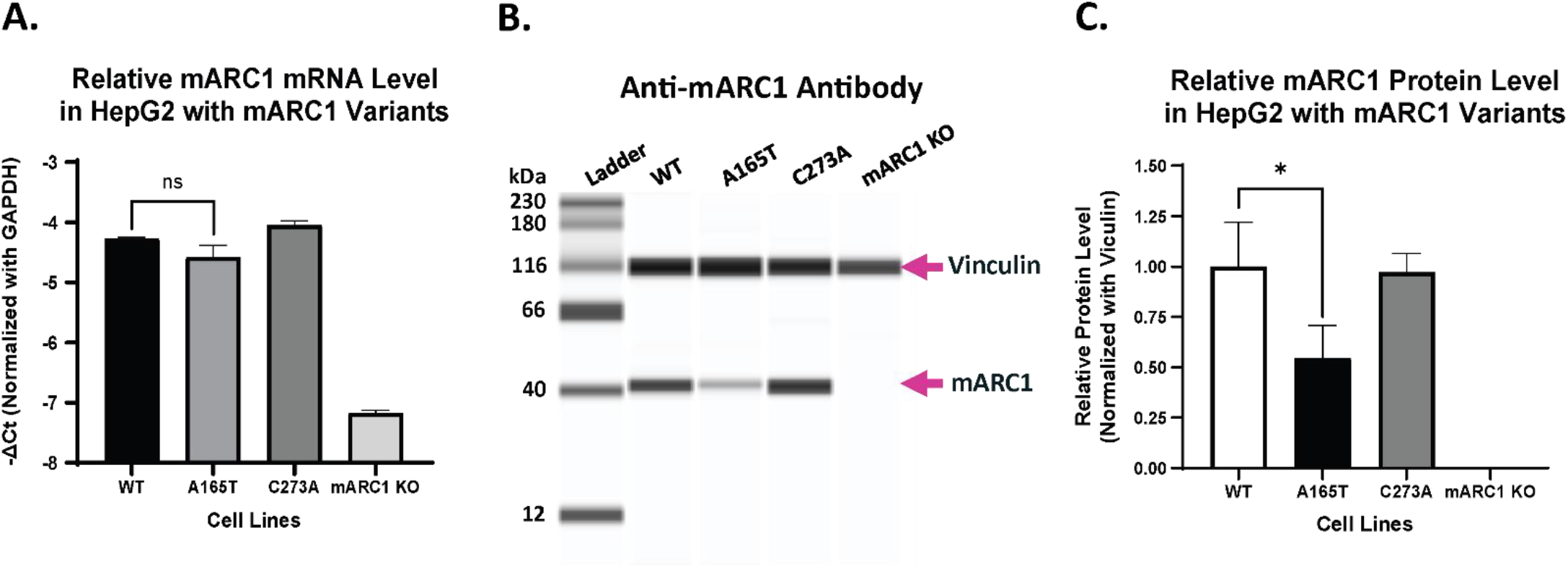
mRNA and protein expression level of mARC1 in HepG2 cells with different genotypes. A. mRNA level in cells of the indicated genotype; B. Jess result for protein expression in the three HepG2 cell lines with MTARC1 WT, A165T and C273A genotype; C. Relative mARC1 protein expression level after normalization (data was first normalized with vinculin level in the same cell line and then normalized with the relative protein level in WT cells) (* P<0.05)

In contrast, genotype-dependent changes in mARC1 protein levels were observed. Significantly lower mARC1 protein levels (∼54%) were detected in the A165T cells relative to WT (Fig. 2B and 2C). mARC1 KO cells did not express mARC1 protein which verifies the specificity of the western analysis. These results indicate that A165T leads to lower levels of mARC1 protein (Fig. 2B & 2C).

### Ectopically expressed mARC1 A165T exhibits reduced stability in double KO cells

A recent study reported that recombinant mARC1 A165T has similar N-reductive activity to WT in a cell-free assay (17,30). This finding may indicate that WT and A165T have similar activities and stabilities outside of a cellular environment. However, our study demonstrates that mARC1 A165T expressed from its endogenous promoter is less stable compared to WT and C273A in HepG2 cells. To further examine the effect of A165T on protein stability, mARC1 variants were ectopically expressed in HepG2 dKO cells using the BacMam system. While no mARC1 protein was detected in the double knockout cells, ectopically expressed mARC1 protein was detected for all genotypes including WT, A165T or C273A. Endogenous mARC1 in WT HepG2 cells served as a positive control and a measure of endogenous expression levels. Notably, the level of mARC1 protein was much lower in the cells expressing A165T variant compared to cells expressing WT or C273A (Fig. 3A). A similar expression pattern was observed in BacMam expressing mARC1-GFP fusion proteins: A165T-GFP showed much lower expression than either WT-GFP or C273A-GFP (Fig. 3B). These results suggest that A165T decreases stability of ectopic mARC1 protein.

**Figure 3.**
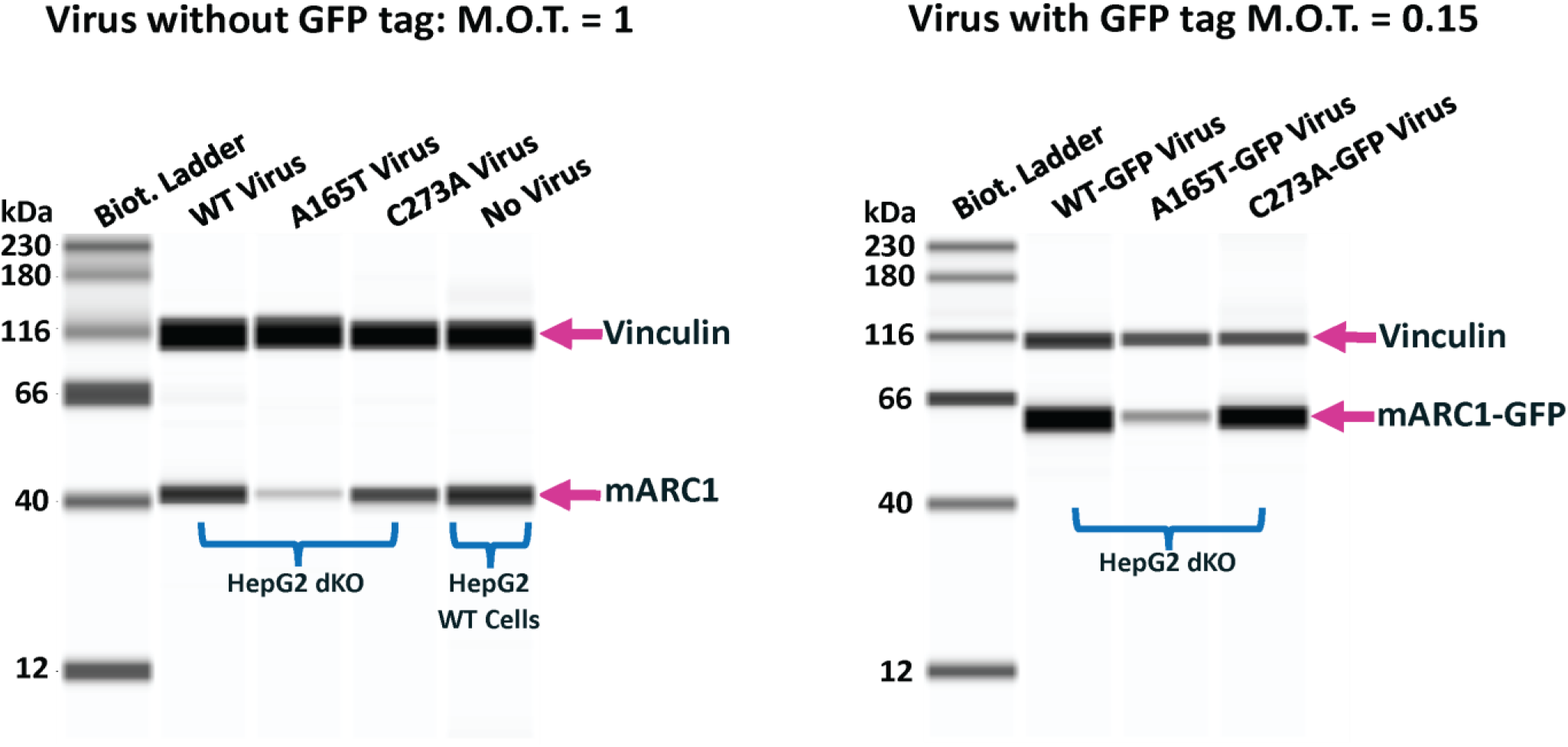
mARC1 protein level in HepG2 dKO cells transduced with BacMam virus expressing mARC1 variants WT, A165T or C273A w/o C-term GFP tag. A. mARC1 protein expression in dKO HepG2 dKO cells transduced with BacMam virus expressing mARC1 variants without GFP tag; B. mARC1 protein expression in dKO HepG2 dKO cells transduced with BacMam virus expressing mARC1 variants with GFP tag.

### Endogenous mARC1 A165T is degraded more rapidly than WT

HepG2 cells expressing mARC1 A165T, either endogenously or ectopically, exhibit lower levels of mARC1 protein compared to cells expressing either WT or C273A. To test whether lower mARC1 A165T levels are due to more rapid protein degradation, HepG2 cells with different *MTARC1* genotypes (WT, A165T and C273A) were treated with two different concentrations (10 μM and 20 μM) of cycloheximide, an inhibitor of protein translation, for 8 hours and mARC1 protein levels were measured. A165T mARC1 protein levels were decreased relative to vehicle at both concentrations of cycloheximide. In contrast, WT and C273A mARC1 protein levels were minimally affected by cycloheximide treatment (Fig. 4).

**Figure 4.**
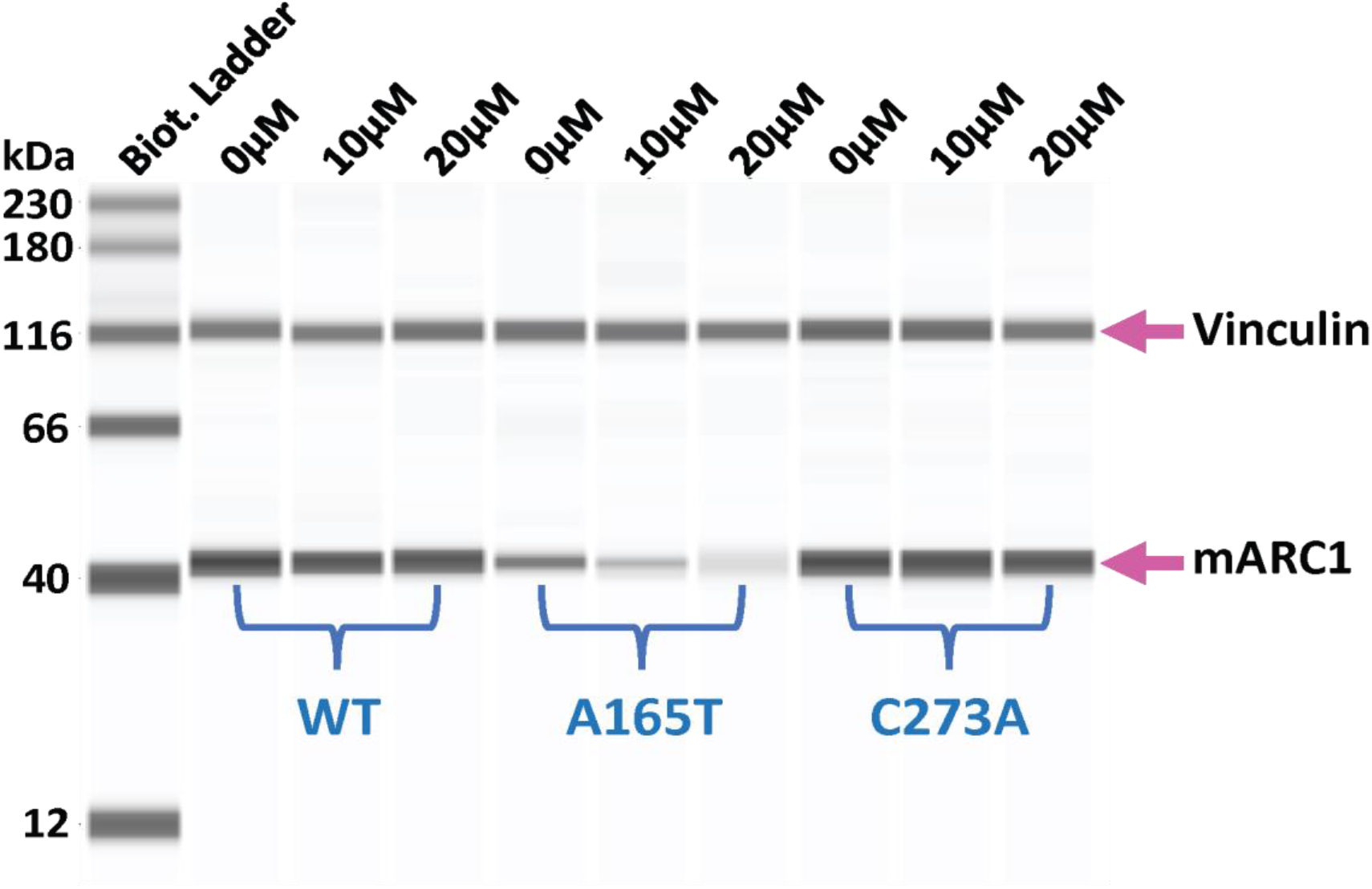
Protein stability analysis for mARC1 WT, A165T and C273A. HepG2 cell lines with three different MTARC1 genotypes WT, A165T or C273A were treated with 0 μM (DMSO only), 10 μM, and 20 μM of cycloheximide for 8 hours. mARC1 protein expression level was examined in those cell lines using ProteinSimple/Jess system. A representative image of 4 repeats of Simple Western results is shown here. (The loading amount of each lane was normalized to vinculin levels to adjust for a small change in vinculin levels with cycloheximide (data not shown))

### Degradation of mARC1 A165T occurs through the ubiquitin-proteasome pathway

To determine whether the ubiquitin-proteasome system is responsible for the rapid degradation of mARC1 A165T, we examined ubiquitination of C-terminal Flag-tagged mARC1 WT, A165T or C273A expressed in HepG2 dKO cells via BacMam transduction. The cells were treated sequentially with cycloheximide and MG-132. Immunoprecipitation (IP) of mARC1-Flag demonstrated that basal ubiquitination of mARC1 was significantly elevated in A165T-expressing cells relative to WT (Fig. 5). Upon treatment with cycloheximide, the ubiquitination signal of mARC1 A165T was reduced to barely detectable (Fig. 5). This effect was less apparent in the WT mARC1-expressing cells (Fig. 5), indicating that WT degradation is slower than the A165T variant. MG-132 treatment prevented the loss of ubiquitinated A165T (Fig. 5), indicating the importance of the proteasome in its degradation. Interestingly, the catalytically dead mutant C273A exhibited a similar ubiquitination pattern as A165T, despite its apparent stability demonstrated in Figure 4.

**Figure 5.**
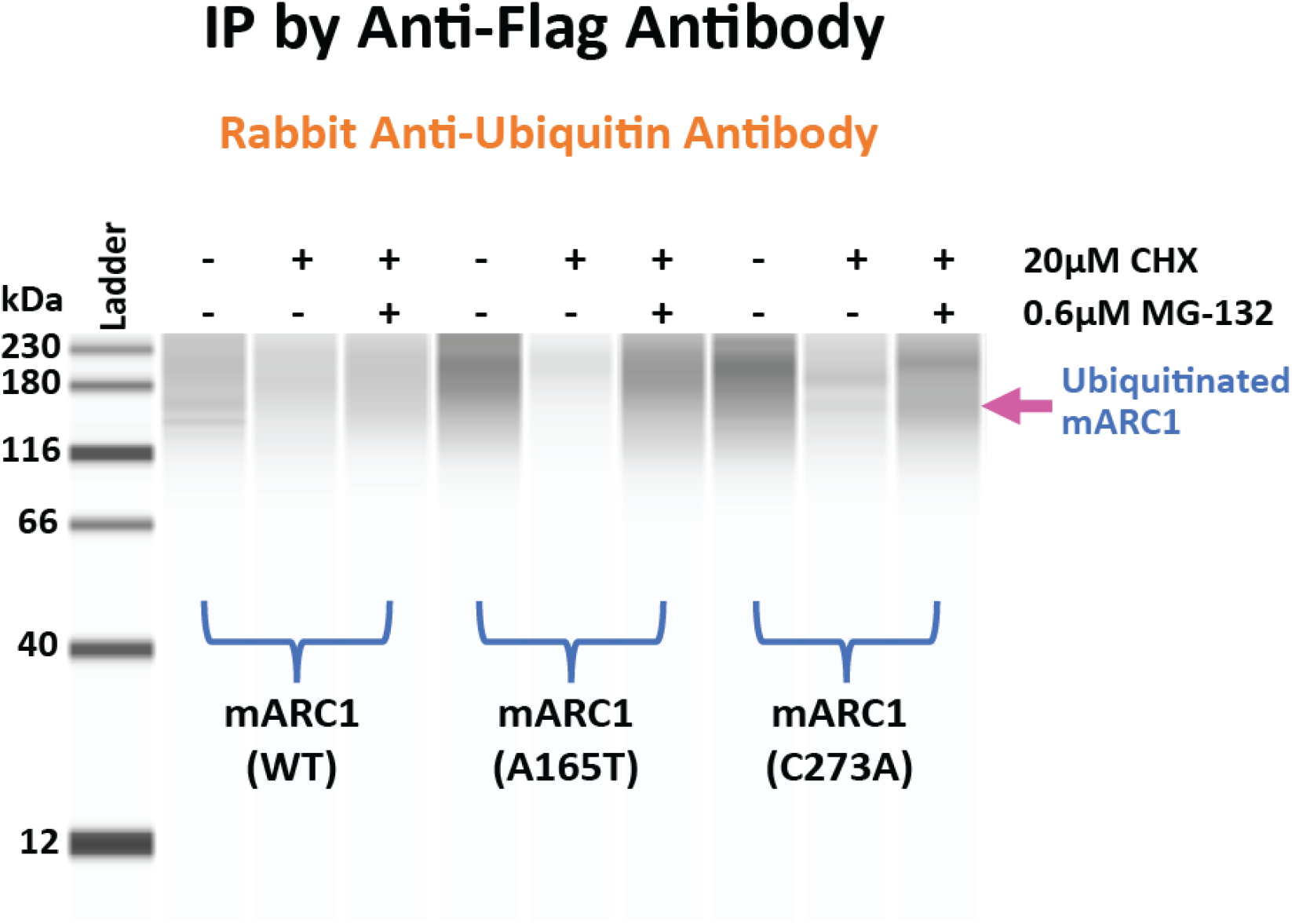
Degradation of mARC1 A165T by ubiquitin proteasome pathway. Western blot showing ubiquitination patterns under different conditions for the three mARC1 variants.

### mARC1 A165T has reduced N-reductive activity in HepG2 cells

The mARC1 enzyme contributes to *in vivo* reduction of N-hydroxylated substrates (19,21). Many studies have demonstrated this catalytic function of mARC1 using recombinant protein (17,20,23,30,32). We designed a cellular substrate turnover assay to evaluate the N-reductive activity of mARC1 variants in HepG2 cells using a model substrate benzamidoxime (BAO) and a putative physiological substrate of mARC1 *N*^ω^-hydroxy-L-arginine (NOHA). We used a flow injection analysis RapidFire-Triple Quadrupole-Mass Spectrometric analysis method (RF-QQQ-MS) to measure the amount of benzamidine (BA) or L-arginine, the products of benzamidoxime and NOHA reduction, respectively, produced by cells and released into the tissue culture media. While HepG2 cells expressing WT mARC1 showed highest efficiency in reducing both substrates, cells expressing the A165T variant exhibited ∼70% of WT reduction capacity in reducing BAO to BA (Fig. 6A) and ∼50% of WT reduction capacity in reducing NOHA to L-arginine (Fig. 6B). This is consistent with the relatively lower protein levels of A165T in this cell line. As expected, cells expressing the catalytically dead mutant C273A had the lowest reduction capacity, exhibiting ∼20% (BAO to BA, Fig. 6A) or 13% (NOHA to L-arginine, Fig. 6B) of WT capacity, which is likely due to the expression of mARC2 protein in these cells. A HepG2 cell line expressing the C273A variant with mARC2 KO showed no reductive activity for either substrate (Supplemental Figure 1).

**Figure 6.**
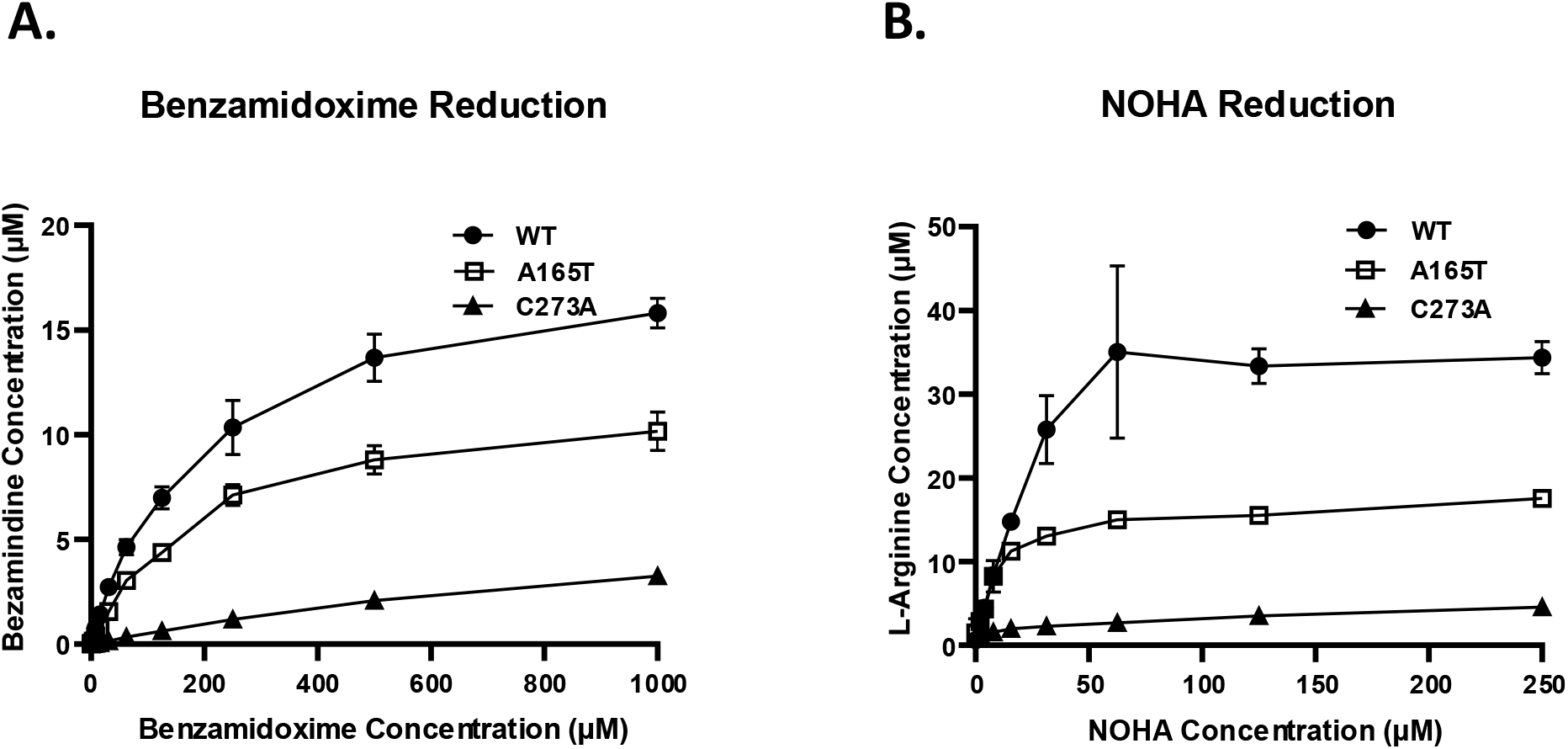
N-reductive activity in HepG2 cells with different MTARC1 genotype: WT, A165T or C273A. A. Benzamidine (BA) generated by the three cell lines in response to the treatment with various concentrations of BAO. B. L-Arginine generated by the three cell lines in response to the treatment with various concentrations of NOHA.

### mARC1 A165T does not affect CYB5B and CYB5R3 expression

The N-reductive activity of mARC1 requires the presence of two additional proteins, cytochrome *b*_5_ type B (CYB5B) and NADH-cytochrome *b*_5_ reductase (CYB5R3) (19,21,22). In our study we examined protein expression level of CYB5B and CYB5R3 in HepG2 cell lines expressing the three mARC1 variants: WT, A165T or C273A. We found high levels of CYB5B and CYB5R3 protein were expressed in all three cell lines that were not remarkably different between genotypes (Fig. 7A and 7B).

**Figure 7.**
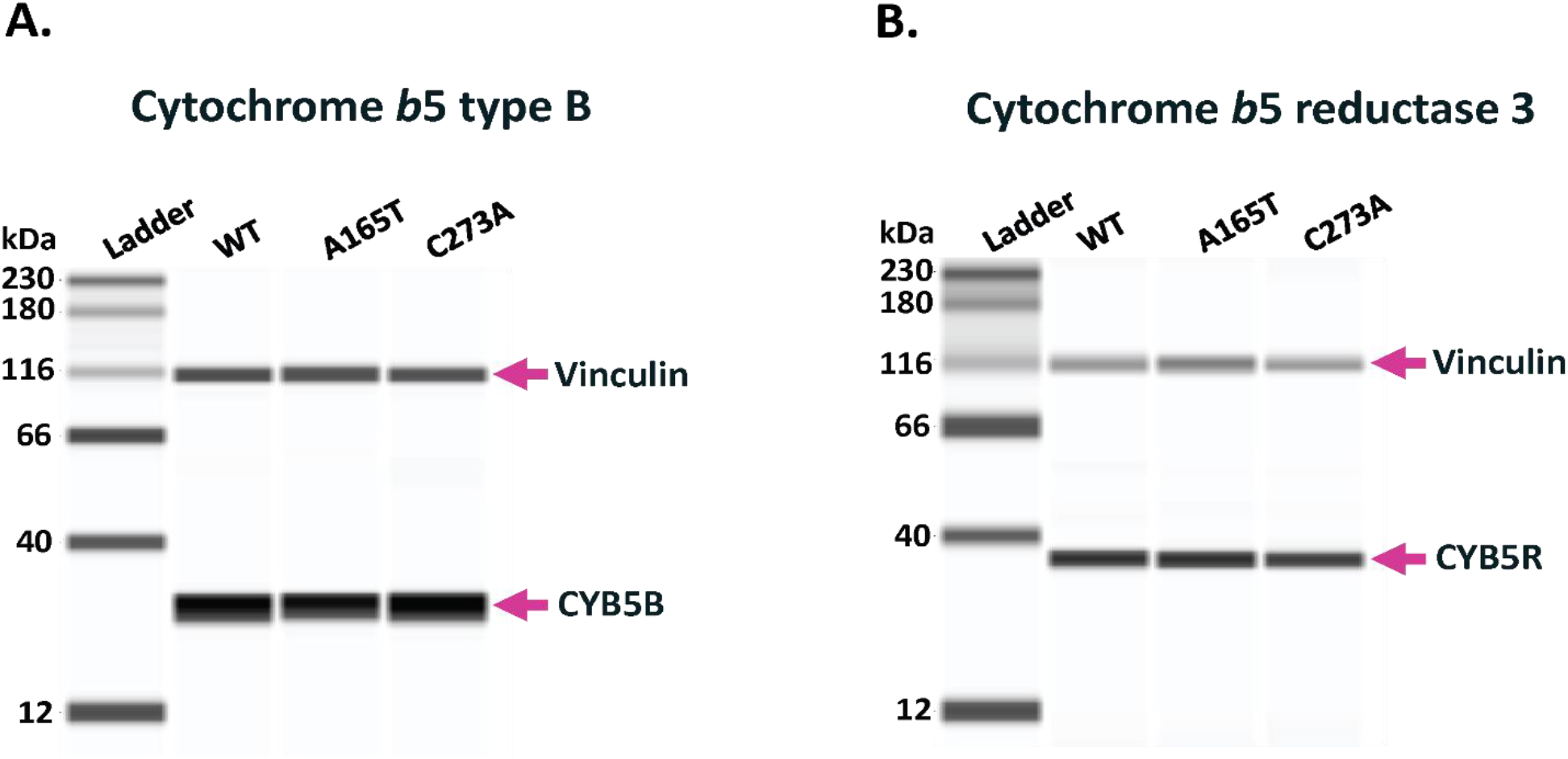
Cytochrome *b*_5_ type B and cytochrome *b*_5_ reductase 3 protein expression in HepG2 cells with different MTARC1 genotypes: WT, A165T or C273A. A. Expression of cytochrome *b*_5_ type B in the three HepG2 cell lines. B. Expression of cytochrome *b*_5_ reductase 3 in the three HepG2 cell lines.

## Discussion and conclusion

NAFLD/NASH is a multifactorial disease. However, recent GWAS have shown that genetic variations play a crucial role in disease susceptibility, development, and severity (1,10,33-37). Some genetic variants such as *PNPLA3* rs738409 C>G (I148M) and *TM6SF2* rs58542926 C>T (E167K) have been found to increase the risk of NAFLD/NASH, while other genetic variants are identified as protective against the disease. *MTARC1* rs2642438 G>A (A165T) is one of recently reported variants which is associated with protection from NASH (1,10,33,36,37). Based on the fact that *MTARC1* p.A165T shows a similar lipid phenotype and protective effect from cirrhosis as the loss of function variant p.R200Ter, it has been surmised that a loss of function induced by A165T may contribute to its association with a decreased risk in NAFLD/NASH (10). However, the underlying mechanism that links *MTARC1* p.A165T variant to its beneficial effect for liver disease remains unclear. Indeed, further characterization of mARC1 A165T protein in hepatocytes, including biochemical and functional properties of the protein, is imperative to obtain a clearer understanding of the biological connection between mARC1 and NASH and provide insight for the development of drug therapies for cirrhosis.

Using CRISPR/Cas9 genome editing, we generated isogenic HepG2 cell lines that express different variants of mARC1, including A165T and C273A. No significant difference in levels of mRNA was found between A165T and WT. Our results are consistent with the published result that mRNA expression level was not significantly affected by *MTARC1* p.A165T in liver tissue samples from patients with NAFLD (11). However, the level of protein expression was largely affected by different genotype. mARC1 A165T showed about 50% reduction of protein expression level when compared to WT protein in HepG2 cells. Moreover, expression of mARC1 A165T in HepG2 cells via BacMam virus transduction at low MOT also showed dramatically less protein compared to WT and C273A when same amount of virus was used. Both experiments demonstrate that the impact of A165T on mARC1 levels occurs at the protein level.

We then further explored the possible cause for the lower protein level of A165T variant and observed that mARC1 A165T showed mislocalization from the outer mitochondrial membrane. mARC1 is known to be localized on outer mitochondria membrane, where the protein functions together with the other two components cytochrome *b*_5_ and cytochrome *b*_5_ reductase to form an intact N-reductive system in the cells (15,38). Subcellular localization of mARC1 is mediated by its N-terminal targeting signal (aa. 1-20) and transmembrane domain (aa.21-40) (15). The C-terminal catalytic Moco-containing domain (aa. 41-337) is exposed to the cytosol (15). Evidence from mARC1 crystal structures and published data showing similar molybdenum content among different recombinant mARC1 variants both demonstrate that residue 165 is not involved in Moco-binding (30,39). A recent crystal structure study of mARC1 A165T showed that there were alternate conformations of Thr165, even though no difference in protein folding and thermal stability was found between A165T and WT (40).

We posit that the alteration of a single amino acid from a hydrophobic to polar amino acid is able to change the anchoring of mARC1 on the outer mitochondrial membrane and cause its mislocalization, even in the absence of significant protein secondary structure changes (41). Our observation that mARC1 A165T is mislocalized to some extent suggests that replacement of Ala by Thr at this position may disrupt mitochondrial anchoring of the protein. Even though this amino acid is not on the transmembrane domain, it could affect interaction with other proteins that might be involved in mARC1 mitochondria membrane targeting and binding (42). Proteins localized on mitochondrial outer membrane have been found to be degraded by ubiquitin-proteasome pathway (43,44). The proteins are polyubiquitinated for degradation in cytosol after they are extracted from the membrane (45,46). When mARC1 protein is detached from the mitochondrial membrane and released into cytosol, it may be more available for polyubiquitination and degradation by cytosolic ubiquitin-proteasome pathway. In our study, we found that A165T more rapidly degraded than the WT and C273A protein when protein translation was blocked with cycloheximide in HepG2 cells. Our study further demonstrated that A165T protein showed greater ubiquitination levels than WT in HepG2 cells. In addition, ubiquitinated A165T degraded faster than ubiquitinated WT, suggesting greater access (i.e. enhanced recognition or entry) to the proteasome. Our data indicate that correct protein localization of mARC1 on the mitochondrial membrane is crucial for maintaining mARC1 protein stability in cells.

In addition to reduced protein level, mislocalization of A165T can result in the loss of function of the mARC1 since it must interact with cytochrome *b*_5_ and cytochrome *b*_5_ reductase on the outer mitochondrial membrane for its N-reductive activity (22). Hudert *et al.* found that there was no change in the level of protein expression of A165T in the liver biopsy sample of NAFLD children compared to wild-type. However, in this same study, a protective effect was still observed for patients carrying the p.A165T variant (13). This apparent discrepancy with our findings may reflect a loss-of-function phenotype that fully manifests from the subcellular mislocalization of A165T without a significant lowering of protein levels in those patient-derived biopsy samples.

After implicating the effect of A165T on mARC1 protein stability and level in cells, we then evaluated the effect of this variant on protein functionality in intact cells using a cell-based substrate turnover assay. In HepG2 cells with A165T genotype, N-reductive activity for both BAO and NOHA was significantly reduced when compared with cells carrying the WT genotype. Previous studies have shown that recombinant mARC1 A165T and mARC1 WT have similar Moco binding efficiency (30) and similar kinetic parameters in reducing sulfamethoxazole hydroxylamine and benzamidoxime (17,30). These data indicate that A165T does not affect N-reductivity of recombinantly expressed mARC1 protein. Taken together with observations from our study, we propose that the decrease in net substrate turnover by A165T in HepG2 cells is due to improperly localized A165T mARC1 protein and separation from its catalytically essential electron transfer partners, with concomitant lowering of total amount of protein.

As expected, the C273A mutation abolished about 80% of N-reductive capability of HepG2 cells. This is consistent with the reported loss of Moco binding by this variant. The residual activity in these cells is likely due to endogenous mARC2 protein, a paralog of mARC1 expressed in these cells which shares similar substrates. The HepG2 cell line expressing the C273A variant with mARC2 KO showed barely any reductive activity, similar to HepG2 dKO cells (Supplemental Figure 1). This is further supported by a rescue experiment in which C273A overexpression was not able to rescue any of the N-reductive activity of the HepG2 dKO cells (data not shown).

We also observed that the expression levels of cytochrome *b*_5_ type B and cytochrome *b*_5_ reductase were not significantly affected by the expression of the A165T variant of mARC1. Even though these proteins are crucial for the N-reductive function of mARC1, it was found that the concentrations of these two proteins were not rate-limiting factors for the N-reductive activity of mARC1 (47). Nevertheless, while our results indicate that the A165T variant does not impact cytochrome *b*_5_ or cytochrome *b*_5_ reductase protein levels, there remains the possibility, beyond the scope of this work, that this variant impacts the efficiency of electron transfer within the reductase complex.

In conclusion, our findings reported here provide biochemical and functional evidence in HepG2 cells supporting the recent GWAS discovery that the A165T variant confers a protective effect on NASH by inducing a functional deficiency of mARC1. Our characterization of subcellular localization and protein stability of mARC1 variants in HepG2 cells support that functional deficiency of the A165T variant in cells is attributable to mislocalization of the variant, which abrogates the interaction of the protein with cytochrome *b*_5_ and cytochrome *b*_5_ reductase for its function. Further, mislocalization of A165T enhanced degradation of the protein by the ubiquitin-proteasome pathway resulting a decreased abundance of the protein (Fig. 8). The results from our study provide a possible explanation why *MTARC1* p.A165T showed similar protective effect from liver disease as the loss-of-function variant p.R200Ter. Together with the fact that the mARC1 protein is involved in lipid metabolism and hepatic fibrotic pathway, our study supports the therapeutic hypothesis that antagonism of mARC1 protein may be a useful method in treating NAFLD and NASH.

**Figure 8.**
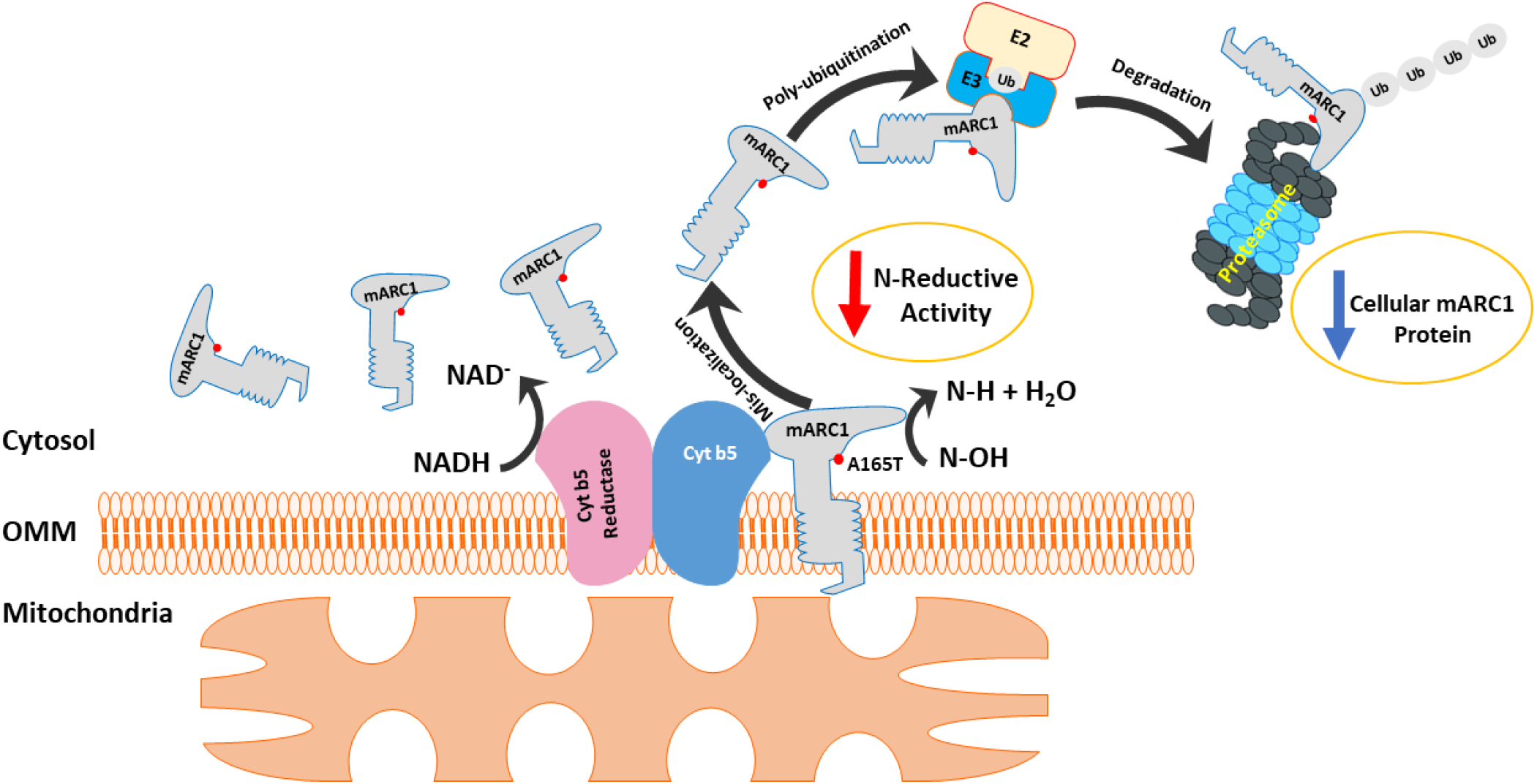
Mislocalization causes fast degradation of mARC1 A165T by ubiquitin/proteasome pathway.

## Experimental procedures

### BacMam virus generation for expressing various mARC1 constructs

Sequence for human *MTARC1* WT, *MTARC1* p.A165T, *MTARC1* p.C273A (reference: NM_022746.4) with or without GFP tag were synthesized and cloned into pHTBV1mcs3, a second generation of BacMam vector (48). (-) control virus was made from pHTBV1mcs3 empty vector. BacMam virus expressing the constructs were then generated via the Bac-to-Bac system. Both plasmid cloning and BacMam virus generation were done by GenScript (Nanjing, China).

### HepG2 cell line culture

HepG2 (ATCC) cells were routinely cultured in EMEM + 10% FBS (ThermoFisher) culture medium and split twice every week.

### HepG2 (*MTARC1*^-/-^, *MTARC*2^-/-^) double knockout (dKO) cell line generation

The guide RNAs (gRNAs) targeting *MTARC1* and *MTARC2* were ordered from Integrated DNA Technologies (IDT). The sequence of gRNA used for *MTARC1* is: GAAGCTGGTTATCCACTGGG; and the sequence of gRNA used for *MTARC2* is: GTAGATCCAGAGCTTCGCCA. Alt-R® *Staphylococcus pyogenes* Cas9 Nuclease V3 and Alt-R® CRISPR-Cas9 tracrRNA were also ordered from IDT. Cas9 gRNA and tracrRNA was first incubated together in tubes to form ribonucleoprotein (RNP) complex. The RNP was then transfected into HepG2 using the 4D Nucleofector system for HepG2 (Lonza). To generate HepG2 dKO cells, HepG2 were first transfected with RNP for *MTARC2* and recovered for 3 days. They were then transfected once again with RNP for *MTARC1*. Three days after the second transfection, cells were plated out into 96-well plates at a density of one cell/well for generating single clonal cell lines. After cells grew to full confluence in each well, the cell lines were screened for *MTARC1* and *MTARC2* genotype via PCR followed by Sanger sequencing. The sequencing results were analyzed using the ICE (Inference of CRISPR Edits) online analysis tool on the Synthego website. Cell lines with KO genotype (frameshift) for both *MTARC1* and *MTARC2* were expanded and pooled together with same cell number from each clonal line. A total of 26 individual clonal cell lines were pooled together for the final double knockout (dKO) cell line used in this study.

### Generation of pooled HepG2 clonal cell lines with *MTARC1* genotype: WT, A165T or C273A

To generate *MTARC1* p.A165T and C273A knock-in (KI) cell lines, RNP targeting *MTARC1* genomic DNA sequence around A165 or C273 were first prepared in tubes. The sequence of gRNA used for targeting around A165 is: GAAGCTGGTTATCCACTGGG; and the sequence of gRNA used for targeting around C273 is: CAGATGCATTTTAACCACAG. ssDNA sequence serving as template for HDR was added to the RNP and then transfected together into HepG2 cells. ssDNA of the template for *MTARC1* p.A165T KI: 5’- GGCTGTGACTTCAGGAAGCTGGTTATCCACTGGGCTGTGGCCTCGCCACAGTCCCTGCCCTCTATCTCCAGGC -3’; ssDNA of the template for *MTARC1* p.C273A KI:5’- TGTCCCCCTTATGATGCTCTGTGTGTGTGTCCAGAGCCATTTTAACCACAGTGGACCCAGACACCGGTGTCATGAGCAGGAAGGAACC -3’. Three days after transfection, cells were plated into a 96-well plate at a density of 1 cell/well for generating single clonal cell lines. Each cell line was then screened for *MTARC1* genotype via PCR followed by Sanger sequencing. The sequencing result was analyzed using the ICE online analysis tool on the Synthego website. For *MTARC1* p.A165T KI, cell lines with correct *MTARC1* p.A165T genotype were picked and expanded. A total of 14 cell lines were pooled together with the same cell number from each clonal line as a final pooled clonal HepG2 *MTARC1* p.A165T cell line used in this study. From the A165T KI experiment, cell lines with *MTARC1* WT genotype were also picked and expanded. A total of 31 cell lines were pooled together with same cell number from each clonal line as a final pooled clonal HepG2 WT cell line used in this study. The same procedure was followed to generate a final pooled clonal HepG2 *MTARC1* p.C273A KI cell line with a mixture of nine single clonal cell lines.

### Subcellular localization of mARC1 protein in HepG2 cells

mARC1 WT, mARC1 A165T and mARC1 C273A BacMam viruses were used to transduce HepG2 dKO cells in suspension (1.2 x 10^5^ cells/ml) at a multiplicity of transduction (MOT) of seven. After the addition of BacMam virus, cells were plated in collagen-coated, 96-well imaging plates (PerkinElmer 6055700) at 1.2 x 10^4^ cells/well in 100 μl media and covered with Breathe-Easy® seals (Sigma Z380059). The plates were incubated in a CO_2_ incubator (37°C, 5% CO_2_) overnight to allow for BacMam protein expression to occur.

On day 2, 100 μl MitoTracker Deep Red (Invitrogen M22426, 100 nM, diluted in culture media) was added to each well of the 96-well plate. The plate was returned to the CO_2_ incubator for 30 minutes. Medium in the plate was gently flicked off and the plate was washed 3x with PBS. Cells in the plate were fixed in 3.7% formaldehyde for 20 minutes and then washed 3x with PBS. The plate was then blocked with 50 μl/well of PERM/BLOCK buffer (HBSS + 0.1% Triton X + 20 mM HEPES + 3% BSA) for 1.5 hr on a slow shaker at room temperature. After the block buffer was gently flicked off, cells were then incubated with 50 μl/well of 1 μg/ml anti-mARC1 antibody (Sigma HPA028702, 1:100 dilution in PERM/BLOCK buffer) and placed on slow shaker overnight at 4°C.

On day 3, the cells were washed 2x with wash buffer (HBSS + 0.1% Triton X + 20 mM HEPES). 50 μl/well of 2 μg/ml secondary goat anti-rabbit AF488 Ab (Invitrogen A32731, 1:1000 dilution in PERM/BLOCK buffer) was added to the plate. The plate was put on a slow shaker at RT for 1 hour and then washed 2x with wash buffer. Hoechst (Invitrogen H3570, 1:3000 in PBS) and CellMask Orange (Invitrogen H32713, 1:20000 in PBS) stains were added to the cells for 30 minutes at RT in the dark. The plate was then washed twice with wash buffer. After the wash buffer was gently flicked off, 100 μl of PBS was added to each well. Cells were then imaged with a PerkinElmer Opera Phenix automated confocal microscope using the 40x water immersion objective. Images were imported to PerkinElmer Columbus 2.9.1 image analysis software for quantification. FITC intensity of mARC1 protein for mitochondrial puncta and background was quantitated for an average from over 6,000 cells (6 wells, each well >1,000 cells)

### PCR and ICE (Inference of CRISPR Edits) analysis for screening the clones with correct genotype

PCR primers for sequence around mARC1 (A165) were: Forward primer: 5’-AAGCATAGCCAGGCCTGTGAATAA-3’; Reverse primer: 5’-TGCAAACTGTAAAAATTCTGGACT-3’. PCR primers for sequence around mARC1 (C273): Forward primer: 5’- AATCTCATCTCAGGGGAATCAACT-3’; Reverse primer: 5’- GTCACATCACTTCACTCCTACAC-3’. PCR primers for sequence for mARC2 KO: Forward primer: 5’- CTGTCTGCCTGTCTTCCTCCATTA-3’, Reverse primer: 5’- TGTCTATGTGTCAGGCCCAAAAGT-3’. PCR products were generated using cell lysate for each cell line as template (DirectPCR lysis reagent, Viagen), and then purified by using QIAquick 96 PCR Purification Kit (Qiagen). The purified PCR products were sent to GENEWIZ for Sanger sequencing. The sequencing results were analyzed for genotype identification with DNASTAR Lasergene software and the Synthego ICE analysis tool.

### Simple Western (Jess) to detect protein expression

The Jess system (ProteinSimple, instrument automated traditional western blotting) was used for protein detection. Cells on plates were harvested with TrypLE™ Select Enzyme (Gibco) and cell pellets were washed with PBS and then lysed with RIPA buffer (ThermoFisher) + Protease Inhibitor (Cell Signaling Technology) + Benzonase Nuclease (Sigma-Aldrich). Jess samples were prepared using the cell lysate following the protocol provided by ProteinSimple. The primary antibody used to detect mARC1 was from Abgent and mARC2 was from Proteintech. An antibody targeting vinculin (Cell Signaling Technology) was always included in detection for each sample as an internal control.

### RT-qPCR to detect mRNA levels of *MTARC1* in various mARC1 KO & KI cells

HepG2 with mARC1 WT, A165T, C273A and dKO pooled clonal cell lines were routinely cultured in T75 flasks. For RT-qPCR experiments, 1 x 10^6^ cells for each cell line were harvested. Following the protocol “TaqMan Gene Expression Cells-to-CT^TM^ Kit” (ThermoFisher), the samples were prepared for RT-qPCR assay. mARC1 assay (ThermoFisher) with the probe targeting 3’-end of mRNA were used in the experiments.

### Benzamidoxime (BAO) substrate turnover assay

HepG2 cells were cultured in 96-well plates with a density of 5.0 x 10^4^ cells/well. 24 hours later, cells were washed with HBSS (STEMCELL technologies) for 10 min. BAO (Sigma-Aldrich) in HBSS at various concentrations (2-fold series dilution starting from 1000 μM) was added to corresponding wells. After cells were incubated with BAO for 3 hours, cell supernatant was harvested for detection of BAO/benzamidine (BA)/3-fluoro-4-methylbenzamidine (FMBA, employed as internal standard (IS)) using RapidFire-Triple Quadrupole-Mass Spectrometric analysis method (RF-QQQ-MS, compromised of RF300 (Agilent Technologies) and Sciex 5000 or 5500 Mass Spectrometer (ABSciex). A graphite C18/type D solid phase extraction (SPE) cartridge (Agilent Technologies) was used for analyte/IS adsorption/elution. The mobile phase A (for sample desalting) was composed of 0.5% (w/v) trifluoroacetic acid in water. The mobile phase B (for elution) was composed of 0.5% trifluoroacetic acid (w/v) in 20% (v/v) acetonitrile/water. Analyte detection was achieved by adsorption/elution: flow rate was 1.5 ml/min, 1.25 ml/min and 1.00 ml/min for pump 1 (mobile phase A), pump 2 (mobile phase B) and pump 3 (mobile phase B), respectively. Multiple Reaction Monitoring (MRM) methodology was used for BAO (Q1 137.0 Da/Q3: 121.0 Da), BA (Q1 121.0 Da/Q3 104.0 Da) and FMBA (Q1 153.0 Da/Q3 136.0 Da) detection.

### *N*^ω^-hydroxy-L-arginine (NOHA) substrate turnover assay

HepG2 cells were cultured in 96-well with a density of 5.0 x 10^4^ cells/well. 16 hours later, cells were washed with HBSS (STEMCELL technologies) for 10 min. NOHA (Sigma-Aldrich) in HBSS at various concentrations (2-fold series dilution starting from 250 μM) was added to corresponding wells. After cells were incubated with NOHA for 3 hours, cell supernatant was harvested for NOHA, L-arginine/^13^C_6_-L-arginine (L-Arg IS) and L-citrulline/^2^H_4_-L-citrulline (L-Cit IS) RF-QQQ-MS detection using RapidFire-Triple Quadrupole-Mass Spectrometric analysis (RF-QQQ-MS) (Sciex 5000/5500 Mass Spectrometry). SPE/mobile phase A/B conditions were the same as those used in the BAO substrate turnover assay. MRM methodology was used for NOHA (Q1 191.1 Da/Q3 146.0 Da), L-Arg (Q1 175.1 Da/Q3 116.0 Da), ^13^C_6_-L-Arg (Q1 181.1 Da/Q3 121.0 Da), L-Cit (Q1 176.2 Da/Q3 159.0 Da) and ^2^H_4_-L-Cit (Q1 180.2 Da/Q3 163.0 Da) detection.

### mARC1 protein stability test in HepG2 cells

HepG2 cells expressing different forms of mARC1 (WT, A165T or C273A) were seeded on 96-well plates with a density of 6.0 x 10^4^ cells/well in 200 μl culture medium and incubated at 37°C in a 5% CO_2_ incubator for 16 hours. A final concentration of 0 μM (DMSO only), 10 μM or 20 μM of cycloheximide (dissolved in DMSO) was added to different wells of each cell line in culture medium (0.0067% DMSO in all wells). After the cells were incubated with cycloheximide for 8 hours, they were then harvested and pelleted. Cell lysates were prepared from the cell pellets and run on ProteinSimple Jess for mARC1 protein detection.

### mARC1 protein immunoprecipitation and ubiquitinated mARC1 protein detection

HepG2 dKO cells were seeded on T150 plates at a density of 1.84 x 10^7^ cells/flask together with one of the BacMam viruses expressing either C-terminal Flag-tagged mARC1 WT, A165T or C273A at an MOT of 1, respectively. Three flasks were prepared for each cell line. Cells were incubated in an CO_2_ incubator for 16 hours. DMSO or cycloheximide (dissolved in DMSO) were added to different flasks with each BacMam virus to achieve a final concentration of 0 μM (DMSO) (1 flask) or 20 μM cycloheximide (2 flasks) in culture medium and incubated with cells for 8 hrs. DMSO or MG-132 was then added to the cells to achieve a final concentration of 0 μM (DMSO) or 0.6 μM MG-132 (1 of the 2 flasks with 20 μM cycloheximide) and incubated with the cells for 2 hrs. A total of three conditions were prepared for cells with each BacMam virus: 0 μM cycloheximide + 0 μM MG-132, 20 μM cycloheximide + 0 μM MG-132, and 20 μM cycloheximide + 0.6 μM MG-132. Cells were then harvested and lysed for immunoprecipitation with Pierce Anti-DYKDDDDK Magnetic Agarose (Invitrogen). For the eluate from the immunoprecipitation, ubiquitinated mARC1 protein was detected with rabbit anti-ubiquitin antibody (Cell Signaling Technology).

### Statistical analysis for all experiments

GraphPad Prism 9 was used for statistical analysis. Four replicates for each condition were performed and outliers were removed if identified (Grubbs’ test, p<0.05). Data comparison was analyzed using unpaired t-test.

**Supplemental Figure 1.**
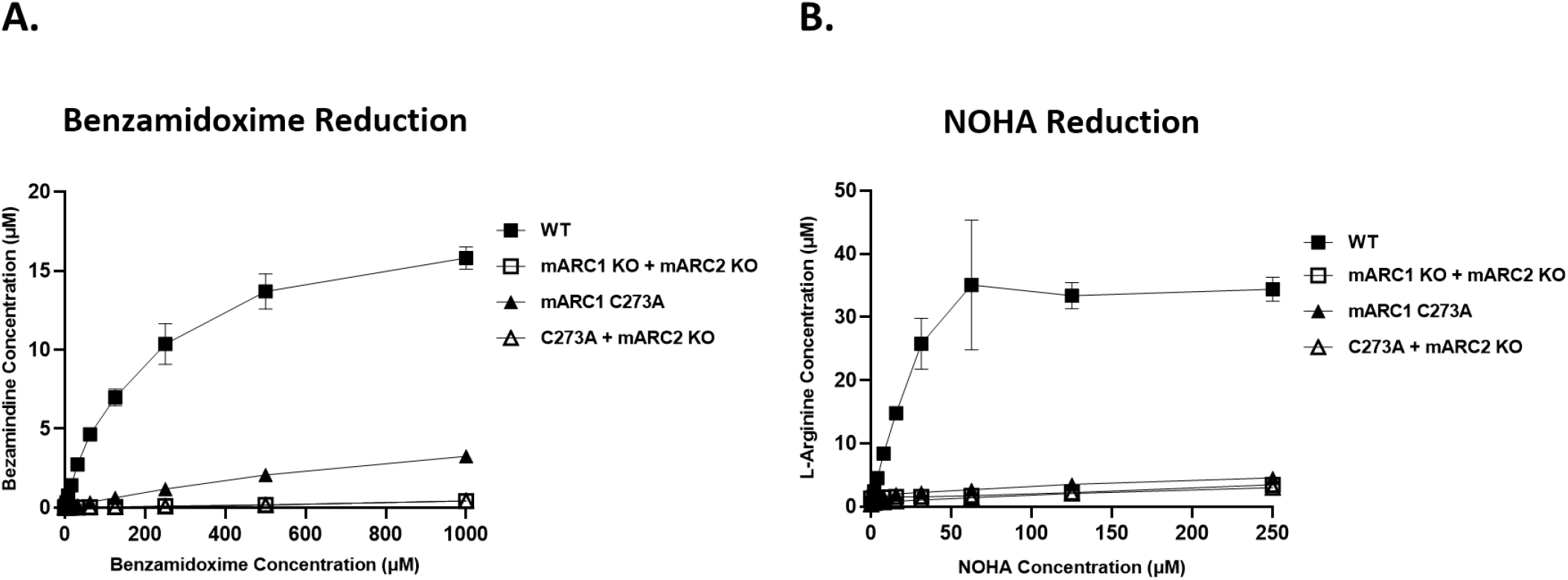
Comparison of N-reductive activity in HepG2 (mARC1 C273A) with HepG2 (mARC1 C273A, MARC2 KO) cells. A. Benzamidine (BA) generated by the four cell lines in response to the treatment with various concentrations of BAO. B. L-Arginine generated by the four cell lines in response to the treatment with various concentrations of NOHA.

## Acknowledgements

We thank the project team for their kind support and helpful discussion, and Mark Harpel for his careful reviews and insightful suggestions.

## Abbreviations

BA: benzamidine
BAO: benzamidoxime
CYB5B: cytochrome b5 type B
CYBR3: cytochrome b3 reductase 3
dKO: double knockout
FMBA: 3-fluoro-4-methylbenzamidine
gRNAs: guide RNAs
GWAS: Genome-wide association studies
ICE: Inference of CRISPR Edits
IS: internal standard
KI: knock-in
KO: knockout
mARC1: mitochondrial amidoxime-reducing component 1
Mo: molybdenum
Moco: molybdenum cofactor
MRM: Multiple Reaction Monitoring
*MTARC1*: Mitochondrial amidoxime-reducing component 1
NAFLD: nonalcoholic fatty liver disease
NASH: nonalcoholic steatohepatitis
NO: nitric oxide
NOHA: *N*^ω^-hydroxy-l-arginine
RF-QQQ-MS: RapidFire-Triple Quadrupole-Mass Spectrometric analysis method
RNP: ribonucleoprotein
ROS: reactive oxygen species
SNPs: single nucleotide polymorphisms
SPE: solid phase extraction
WT: wild-type

